# Autoantibodies against eukaryotic translation elongation factor 1 delta (EEF1D) in two patients with autoimmune cerebellar ataxia

**DOI:** 10.1101/2023.09.05.556333

**Authors:** Liyuan Guo, Haitao Ren, Siyuan Fan, Xingchen Chao, Mange Liu, Hongzhi Guan, Jing Wang

## Abstract

To report a novel autoantibody against eukaryotic translation elongation factor 1 delta (EEF1D) in two patients with autoimmune cerebellar ataxia (ACA). The patients on one center with cerebellar ataxia of unknown cause, who were tested positive with tissue-based indirect immunofluorescence assay (TBA) on rat cerebellum sections and negative for comprehensive anti-neural autoantibodies panel, were investigated for novel autoantibody identification. Tissue-immunoprecipitation (TIP) combined with mass spectrometric (MS) analysis was used to identify the target antigen. The EEF1D protein was identified from complex precipitated by serum and cerebrospinal fluid (CSF) of one patient, while the specific binding of autoantibodies and EEF1D were confirmed by subsequent neutralization experiment, recombinant cell-based indirect immunofluorescence assay (CBA) and western blot. In consequently test using CBA, another anti-EEF1D positive ACA patient was identified and confirmed by western blot. All enrolled 30 health doners and 15 connective tissue diseases patients without neurological disorders were anti-EEF1D negative. The two anti-EEF1D positive patients presented similar clinical manifestations, their outcomes supported the effectiveness of immunotherapy, but the cerebellar atrophy occurred before treatment may be irreversible. Symptoms of one patient worsened again after the weaning of pulse corticosteroid therapy. The identification of anti-EEF1D autoantibody provides a new potential biomarker for early diagnosis and recurrence prevention of ACA.

## 1 Introduction

Autoimmune cerebellar ataxia (ACA), which is an important cause of sporadic cerebellar syndrome and can lead to severe disability, is a potentially treatable disorder that may recover with immunotherapy (1). Autoantibodies are useful biomarkers for the early detection and diagnosis of ACA. While a considerable number of ACA related autoantigens have been identified, there are still many ACA patients whose serological profiles cannot match to any of classified antigen. This limits the diagnosis and treatment. In current study, we detected novel autoantibodies from two ACA patients and identified eukaryotic translation elongation factor 1 delta (EEF1D) as the target antigen.

## 2 Samples and methods

### 2.1 Samples

Patients with cerebellar ataxia of unknown cause who were registered to the program of encephalitis and paraneoplastic syndrome (PNS) of Peking Union Medical College Hospital (PUMCH) from July 2018 to February 2023 were enrolled. This study was approved by the Ethics Committee of PUMCH (JS-891 and JS-2184), and informed consent was obtained from each patient. Sera of enrolled patients were first tested using a tissue-based indirect immunofluorescence assay (TBA) on rat cerebellum sections. Positive sera were further screened for well-established anti-neural autoantibodies using recombinant protein by immunoblots for intracellular or cell-based assays (CBAs) for extracellular autoantibodies (including antibodies target to MOG, AQP4, NMDA-R, CASPR2, AMPA-R, LGI1, GABAb-R, GAD-65, ITPR1, ZIC4, PKCγ, AP3B2, PCA-2, CARP VIII, Homer-3, NCDN, CV2/CRMP5, PNMA2, Ri, Yo, Hu and amphiphysin), commercial test kits were purchased from Euroimmune, Germany. When a TBA-positive serum was negative for the above autoantibody screenings, the sample was investigated for the identification of a new antibody. Additionally, 15 age- and sex-matched systematic autoimmune disease patients without neurological manifestations and 30 health doners were enrolled.

### 2.2 Tissue-based indirect immunofluorescence assay (TBA)

The TBA was performed as described by Guo *et al*. (2). Each slide with rat cerebellum section was incubated with 30 μl of sample diluted in phosphate-buffered saline (PBS) at 4 °C for 3 h, and then incubated with fluorescein isothiocyanate (FITC) labeled polyclonal goat anti-human IgG (Cat. ZF-0308, ZSBio, China) at room temperature for 30 min. For the colocalization test, slides were incubated with 30 μl of sample diluted in PBS (1:100) and anti-EEF1D rabbit antibody (Cat. Ab153728, Abcam, USA) at 4 °C for 3 h, then FITC labeled goat anti-human IgG (Cat. ZF-0308, ZSBio, China) and Alexa Fluor 555 labeled goat anti-rabbit IgG antibody (Cat. Ab150078, Abcam, UK) were co-incubated at room temperature for 30 min. For evaluation of IgG subclasses, patient serum was tested on rat cerebellum sections as described above, with the following modifications: unconjugated sheep anti-human IgG antibodies specific for IgG subclasses 1 to 4 (Nodics-Mubio, Netherlands, 1:100) were substituted for the FITC labeled goat anti-human IgG antibody, and AF568 labeled donkey anti-sheep IgG (Invitrogen; absorbed against human IgG, 1:200) was used to detect the subclass specific antibodies. TBA results were examined by two independent observers using a DMi8 microscope (Leica, Germany).

### 2.3 Tissue-immunoprecipitation (TIP) and Mass spectrometry (MS)

TIP was performed as described by Scharf *et al*. (3). Rat cerebellum wase homogenized after shock-frozen in liquid nitrogen, the tissue lysates was centrifuged at 13,000 × g and 4 °C for 10 min, supernatants were incubated with serum (1:33 diluted) or cerebrospinal fluid (CSF) (undiluted) at 4 °C for 3 h. Immunocomplexes were precipitated with Protein G Dynabeads (Cat. 10003D, Thermo Fisher Scientific, USA) at 4 °C overnight, and then eluted with 60 µl lysis buffer with Laemmli sample buffer (Cat. S3401, SIGMA, USA). The eluted proteins were analyzed by using an Easy nLC 1000 (Thermo Fisher Scientific,USA) coupled to a Q-Exactive Plus mass spectrometer (Thermo Fisher Scientific,USA), as described by Guo *et al*. (2). Raw MS data were analyzed using Proteome Discoverer 2.0 software (Thermo Fisher Scientific, USA) to identify the proteins and searched against *Rattus norvegicus* proteins from the UniProt sequence database. Protein identification was supported by at least one unique peptide. The results were filtered based on a false discovery rate (FDR) of less than 1%.

### 2.4 Recombinant expression of full-length EEF1D in HEK293 Cells

HEK293 cells (CRL-1573, ATCC, VA, USA) were cultured in high-glucose DMEM (Cat. No. 11965092, Life Technologies, USA) supplemented with 10% FBS at 37 °C and 5% CO_2_. The pcDNA3.1 plasmid inserted full-length human EEF1D cDNA (NM_001130053) was customized purchased from GENEWIZ, China, a 6X His-tag was added at the C-terminal. Transient transfection was performed by using FuGene HD transfection reagent (Cat. E2311, Promega, USA). For CBA, cells were seeded on sterile cover glasses 24 hours before transfection, 48 hours after transfection cells were fixed with 4% PFA. For western blot and neutralization experiments, transient transfection was performed in the same way, and total protein of transfected cells was obtained by lysis buffer (Cat. P0013D, Beyotime, China) 48 h later. The cell lysate was centrifuged at 13,000 × g and 4 °C for 10 min, and the recombined EEF1D proteins were purified from supernatant by Ni^2+^ affinity chromatography.

### 2.5 Western blot analysis

Purified EEF1D proteins were electrophoresed on 10% SDS-PAGE gels, transferred to PVDF membranes, blocked for 1 h in 5% nonfat milk, and incubated with serum from patients or health doners (1:100) or anti-EEF1D rabbit antibody (Cat. Ab153728, Abcam, USA) overnight at 4 °C respectively. Detection of the primary antibodies was performed with a 1:5000 diluted HRP-conjugated anti-human or anti-rabbit IgG (Cat. A0201 and A0208, Beyotime, China) for 60 min, and detected using an enhanced chemiluminescence (ECL) kit (Cat. CW0049, CWBio, China).

### 2.6 Neutralization experiments

Patient’s serum was 1:1600 diluted and preincubated with 1 ug purified EEF1D protein at 4 °C overnight, and then TBA was performed.

### 2.7 Recombinant cell-based indirect immunofluorescence assay (CBA)

Recombinant HEK293 cells overexpressing his-tagged human full-length EEF1D and mock-transfected control cells were incubated with serum or CSF and anti-His tag mouse antibody (Cat. AH367, Beyotime, China) for 2 h at room temperature. The binding of human IgG was detected by FITC labeled goat anti-human IgG antibodies (Cat. ZF-0308, ZSBio, China). Alexa Fluor 555 labeled goat anti-mouse IgG antibodies (Cat. Ab150114, Abcam, USA) and DAPI (Cat. C1002, Beyotime, China) were also added to present EEF1D protein and cell nuclei. The slides were observed using a DMi8 microscope (Leica, Germany).

## 3. Results

### 3.1 Identification of the novel autoantigen EEF1D

From July 2018 to November 2019, we recruited five ACA patients whose serum reacted with rat cerebellum but not with established neural autoantigens, one of them was identified as anti-Rab6a/b positive (2). Sera and CSF of other four patients, including the first patient in current study, were involved for further autoantigen identification. Serum and CSF of the index patient have a specific IgG immunostaining pattern in neuronal soma staining in Purkinje cell layer and molecular layer of rat cerebellum (Figure 1A). The IgG subclass repertoire of the autoantibody includes IgG1, IgG2, and IgG3 (Figure 1B).

**Figure 1.**
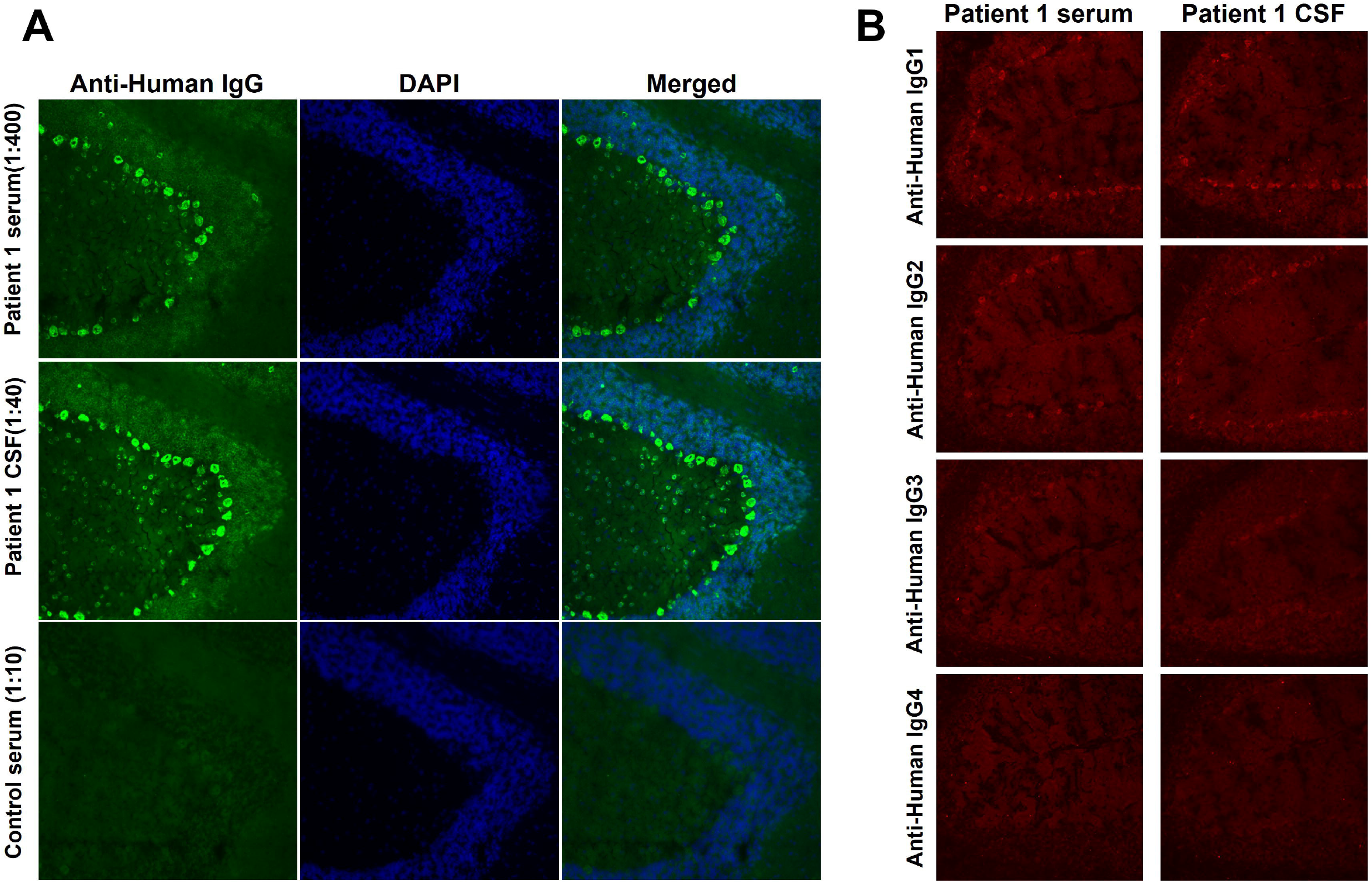
IgG immunofluorescence straining in rat cerebellum. A: Cryosections of rat cerebellum were incubated with patient 1 serum (1:400 diluted) and CSF (1:40 diluted) or control serum (1:10 diluted) in the first step and with FITC-labeled goat anti-human IgG in the second step (green). Nuclei were counterstained by incubation with DAPI (blue). A fine IgG staining was obtained in in both Purkinje cell layer and molecular cell layer. B: Subclass analysis revealed that the autoantibodies belong to IgG1, IgG2, IgG3, but not IgG4.

Immunocomplexes of rat cerebellum and the index patient serum were analyzed by MS. The protein EEF1D was identified as candidate autoantigen since it didn’t appear in immunocomplexes in parallel precipitated by sera from health doners or systematic autoimmune disease patients. As shown in Figure 2, homology of rat Eef1d protein and human EEF1D is high. Human EEF1D gene encodes different spliced protein isoforms, the long isoform (NP_115754) is only expressed in brain and testis, and the shorter one (NP_001951) is ubiquitously expressed. In rat, there are isoforms of different lengths too (Uniprot ID: Q68FR9-1 and Q68FR9-2). The Proteome Discoverer 2.0 software utilized Q68FR9-1 (the shorter one) to analyzed peptide by default. As a result, 17 peptides were identified in EEF1D with more than 60% protein coverage (adjacent or overlapping peptides are merged in Figure 2).

**Figure 2.**
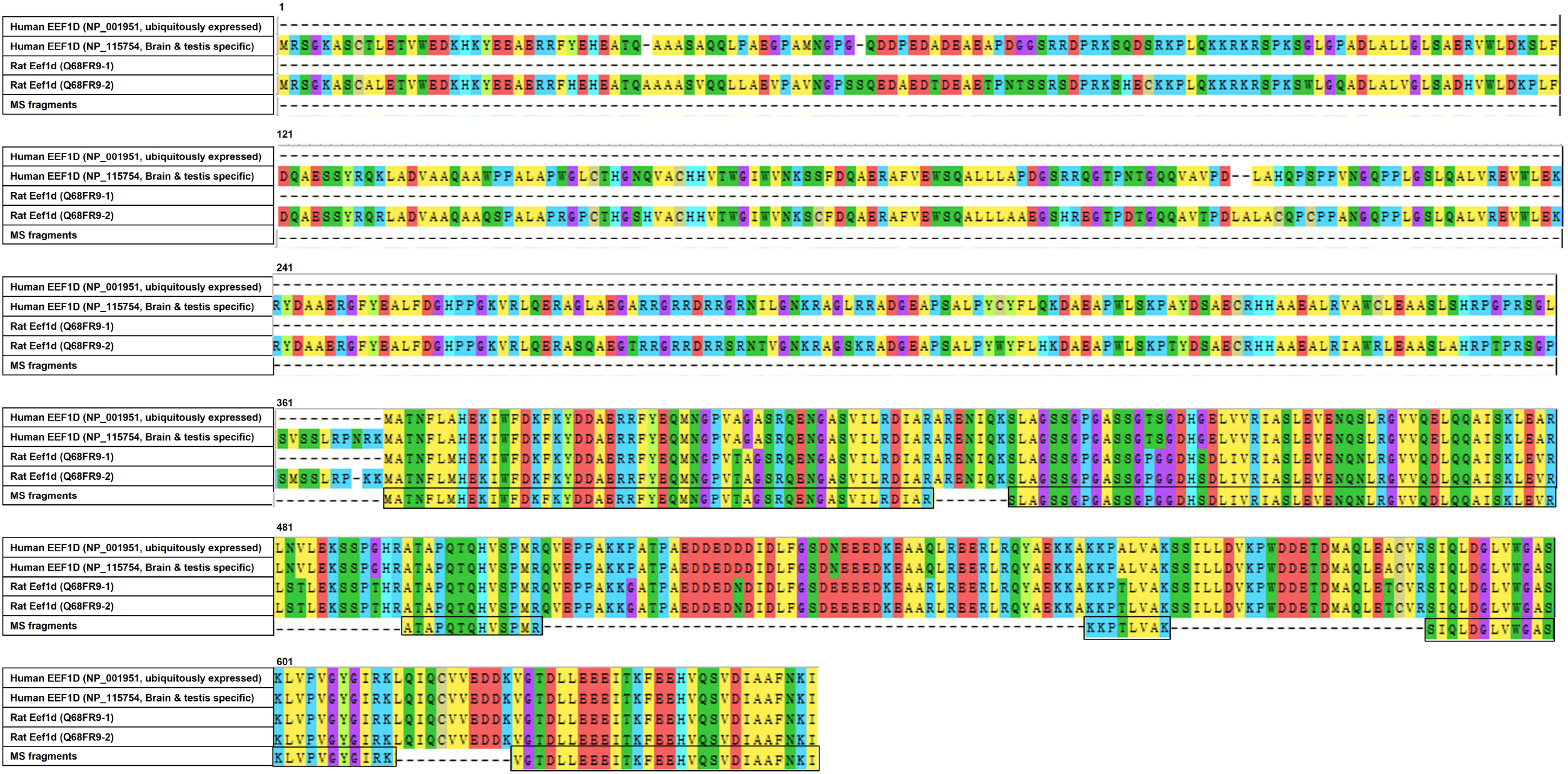
Sequence alignment between MS fragments and human/rat EEF1D protein. Peptides sequences was aligned with both long and short isoforms of human/rat EEF1D protein.

Considering the brain specific expression of long isoform, we recombinant expressed the full-length EEF1D cDNA for further detection. From August 2020 to February 2023, 33 other ACA patients without clear serological profiles were enrolled in. Among them, the second anti-EEF1D IgG positive patient was detected. As shown in Figure 3A, specific binding of EEF1D protein in western blot with sera of the two patients was detected, not in with sera of health controls. In TBA on rat cerebellum, anti-EEF1D antibody preformed neuronal soma staining, similar to serum and CSF of the index patient (Figure 3B). After neutralizing with human EEF1D protein, the staining of patient samples was abolished in TBA on rat cerebellum (Figure 3C).

**Figure 3.**
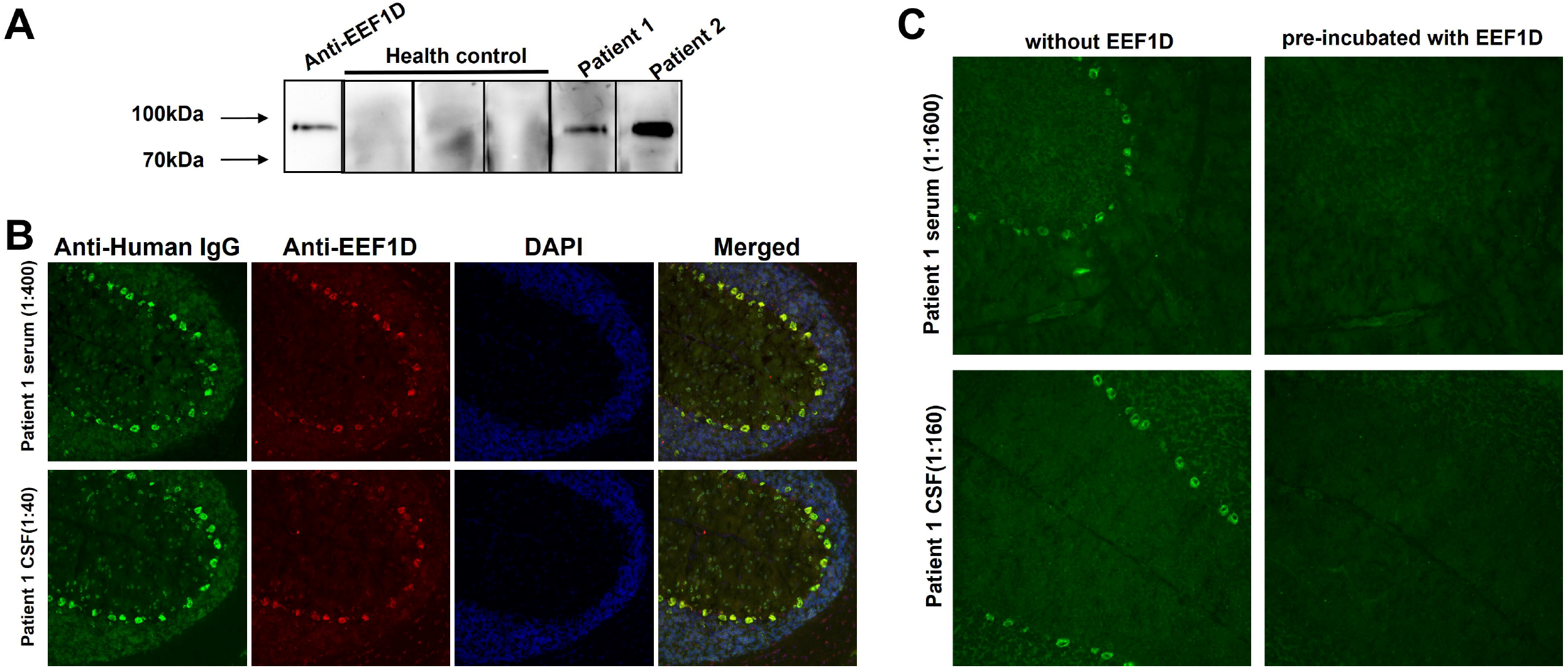
Specific binding between autoantibodies and human full length EEF1D. A: Western blotting shows reactivity of serum samples with recombinant human EEF1D protein in two ACA patients, not in health controls. B: Immunofluorescence staining of rat cerebellum tissue section with patient’s serum (green) and anti-EEF1D(red) antibody. The merged images (right, yellow) show localization of the reactivity in the same region. Nuclei were counterstained by incubation with DAPI (blue). C: After pre-incubating with recombinant human EEF1D protein, the staining of patient samples was abolished.

### 3.2 Specificity of anti-EEF1D autoantibody

Sera from 15 systematic autoimmune disease patients without neurological manifestations and 30 health doners were analyzed by CBA in parallel to the sample of the index patients. None of these control sera produced positive immunofluorescence pattern on the recombinant EEF1D substrate. As shown in Figure 4, sera and CSF of the two patients showed positive staining, which is specifically distinguished from sera of health control. This result suggested that the anti-EEF1D autoantibody was specific to disease of ACA, but not in systematic autoimmune disease patients or health people.

**Figure 4.**
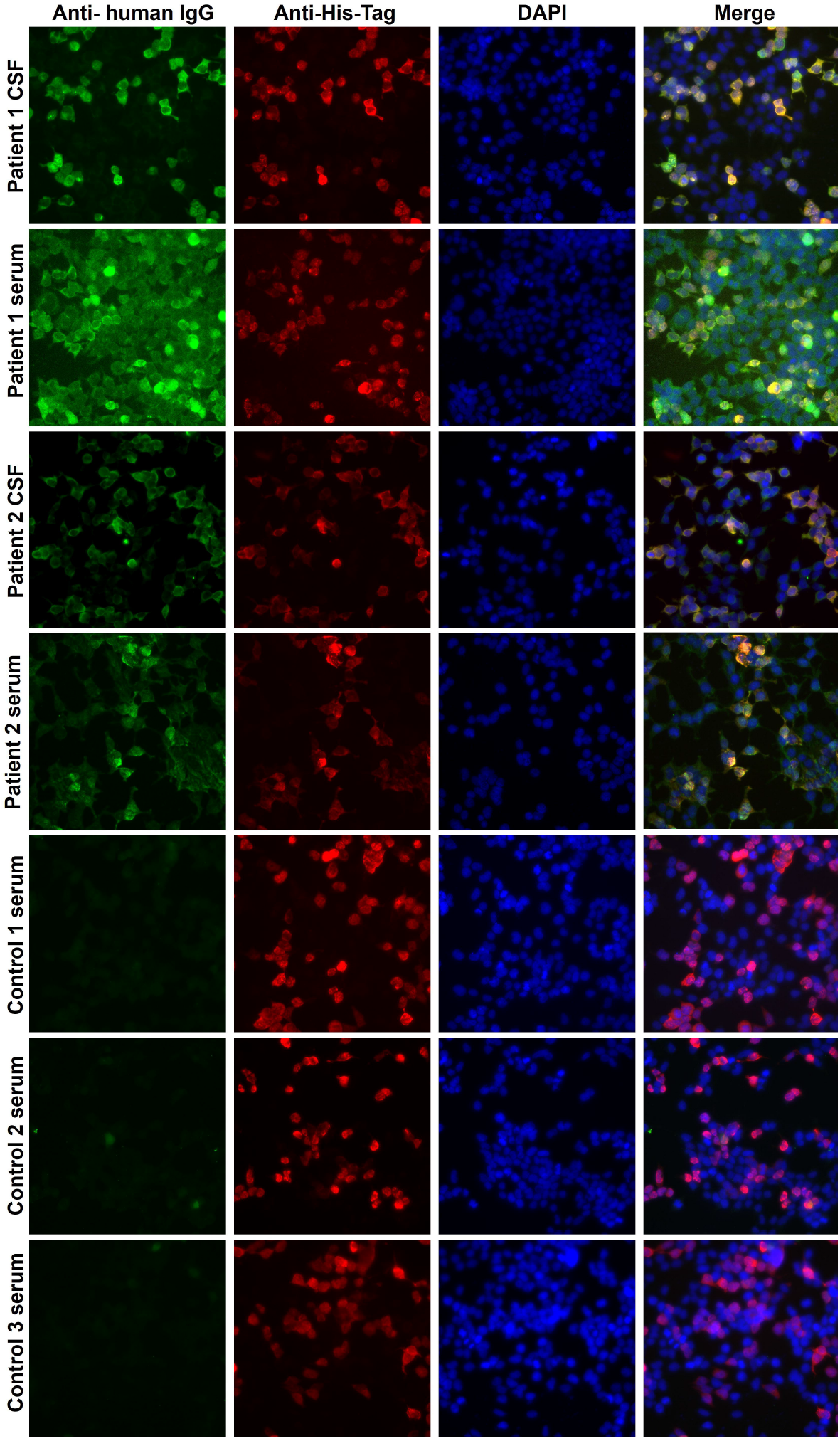
Distinguished CBA patterns of samples from anti-EEF1D positive patients and health controls. Formaldehyde-fixed recombinant HEK293 cells expressing human EEF1D protein were incubated with samples from index patients or health controls. None of control sera produced a similar immunofluorescence pattern to that of the patient samples on the recombinant EEF1D substrate.

### 3.3 Clinical data of anti-EEF1D positive patients

#### Case 1

The 43-year-old female presented in June 2019 with progressive dizziness, tinnitus and unstable gait for 3 months. Her past medical history was noted for connective tissue disease for which she was taking mycophenolate mofetil (MMF) and low-dose corticosteroid. Physical examination showed a stumbling gait, unsteadiness in heel-to-shin test and decreased tendon reflexes in left upper extremity. There was no dysarthria, nystagmus or pathological reflex. CSF white blood cell count was 4/μL while CSF-specific oligoclonal bands (SOBs) were positive. She was also positive for anti-SSA and anti-nuclear antibodies. Brain magnetic resonance imaging (MRI) performed in May 2019 showed multiple hyperintensity lesions in DWI and FLAIR sequences, which disappeared 1 month later (Figure 5). Neuroconductive study revealed decreased sensory nerve action potential in bilateral median nerves and right ulnar nerve. Screening for infection, toxins, metabolic diseases and malignancy was unremarkable. ACA with peripheral neuropathy was diagnosed and she received pulse corticosteroid therapy. MMF treatment was maintained. Her symptoms transiently improved but then continued to deteriorate, and she complained of episodic clonic shaking of the left extremities. Brain MRI conducted in October 2019 revealed cerebellar atrophy. In the video follow-up in January 2023, she was bedridden and the Scale for Assessment and Rating of Ataxia score was 23.

**Figure 5.**
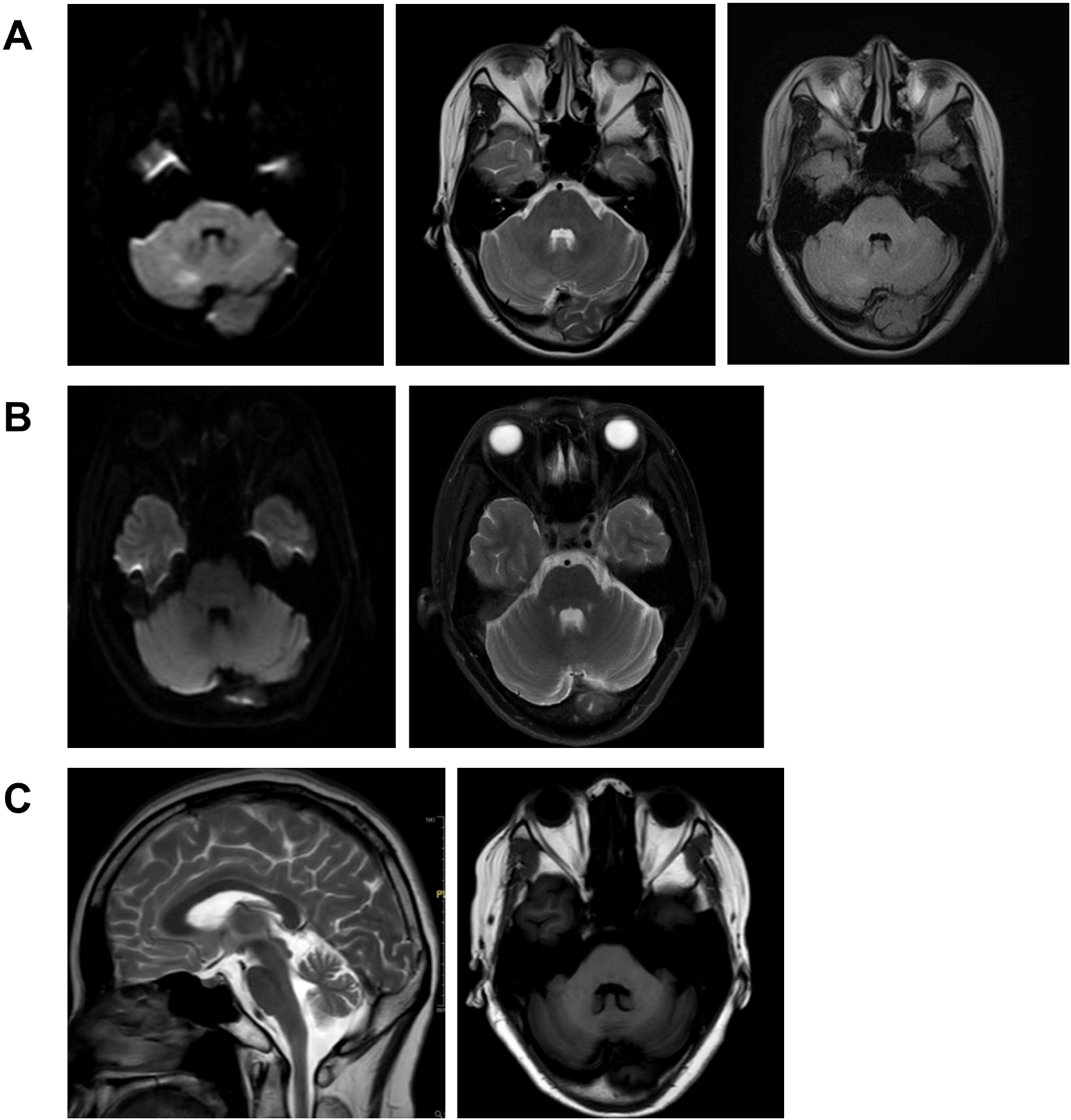
Brain magnetic resonance imaging of anti-EEF1D positive patient #1. A: Brain MRI scans conducted two months after disease onset (May 2019) showed multiple hyperintense lesions in the cerebellum of patient #1. B: One month later (June 2019), follow-up brain MRI scans showed that the lesions in the cerebellum of patient #1 had disappeared. C. MRI conducted in October 2019 showed cerebellar atrophy.

#### Case 2

The 59-year-old female presented with unstable gait for 5 years and paresthesia in bilateral calve for 2 weeks. Physical examination revealed gait and bilateral limb ataxia with decreased tendon reflex in lower extremities. Sensation was intact. She was positive for anti-SSA and anti-nuclear antibodies. CSF examinations were unremarkable except for the presence of SOBs. Cerebellar atrophy was noted in brain MRI. Malignancy screening was negative. She was diagnosed with ACA and peripheral neuropathy. After pulse corticosteroid therapy, intravenous immunoglobulin and MMF treatment, her symptom stopped alleviating and remained stable for 3 years. Clinical data of the two patients were summarized in Table 1.

**Table 1.** Clinical findings, therapies, and outcome of two anti-EEF1D IgG positive patients.

| Patients #,<br>sex, age at<br>al onset | Clinical<br>summary | Systemic<br>autoantibodie<br>s/<br>autoimmune<br>diseases | Brain MRI | CSF findings | Immunothe<br>rapy | Follow<br>-up<br>time<br>(month<br>s) | Outcome |
| --- | --- | --- | --- | --- | --- | --- | --- |
| #1<br>Female, 43 | Subacute<br>cerebellar<br>syndrome<br>with<br>peripheral<br>neuropathy | Anti-SSA,<br>ANA,<br>connective<br>tissue disease | Multiple lesions<br>in the<br>cerebellum<br>which disappear<br>1 month later,<br>cerebellar<br>atrophy in<br>follow-up | WBC 4/μL,<br>SOB (+) | Corticoster<br>oids, MMF | 40 | Worsened<br>after transient<br>improvement |
| #2<br>female, 54 | Chronic<br>cerebellar<br>syndrome<br>with<br>peripheral<br>neuropathy | anti-SSA,<br>ANA | Cerebellar<br>atrophy | WBC 2/μL,<br>SOB (+) | IVIg,<br>corticostero<br>ids, MMF | 33 | Stabilized |

ANA: anti-nuclear antibody; CSF: cerebrospinal fluid; IVIg: intravenous immunoglobulin; MMF: mycophenolate mofetil; OB: oligoclonal bands; WBC: white blood cell.

## 4 Discussion

Up to now, about 30 types of ACA related anti-cerebellar autoantibodies have been reported (4). Although most ACA-related autoantibodies are targeted to intracellular compositions and are regarded non-pathogenetic (5), they serve as important clues for immune-mediated mechanism, since they are specific for ACA and not detected in serum or CSF samples of patients with other systematic autoimmune disease, with ataxia of other etiology or healthy subjects. Serological detections that target specific autoantigens can aid the diagnosis and treatment for patients with unexplained cerebellar ataxia symptoms.

In the current study, we identified anti-EEF1D IgG antibodies in two ACA patients. The diagnosis was based on the inflammatory CSF profile, the presence of systematic autoantibodies and the exclusion of other etiologies, according to the diagnostic criteria of the International Task Force (1). The subclasses of autoantibodies include IgG1, IgG2, and IgG3, similar to previously identified anti-neural antibodies (6).

EEF1D is a protein subunit of the eukaryotic translation elongation 1 complex, which is localized intracellularly and plays essential roles in aminoacyl-tRNA related protein translation elongation (7, 8). Human EEF1D gene encodes at least four different splice isoforms, the longest one (647 aa) is spatial-specifically expressed in brain and testis, and the other shorter ones (252-281 aa) are ubiquitously expressed (9). Mutations in EEF1D gene have been identified as causes of several neurodevelopmental disorders (9). Deletion of the long isoform in mice leads to seizures and abnormal behaviors (10). EEF1D has also been reported overexpression in various tumors, such as glioblastoma, glioma, lymphoma (11-14). Additionally, there are eight pseudogenes of EEF1D in human genome, EEF1D pseudogene-derived proteins or short peptides were reported correlated to a range of tumors and neurological diseases, such as breast carcinoma, ankylosing spondylitis, non-small cell lung cancer, lymphoma (15).

Since the EEF1D protein is an intracellular antigen, the pathogenic roles of autoantibodies against EEF1D needs further study. Outcomes of these two patients supported the effectiveness of immunotherapy. One patient had a brief remission of symptoms but worsened again soon after the weaning of pulse corticosteroid therapy, suggesting the need for more intense and durable immunosuppressive treatment. Another patient stabilized after immunotherapy, which indicated that the aberrant autoimmune process might been suppressed. But lost cerebellar function, as implicated by the cerebellar atrophy, might not be reversed. Therefore, it is important to start immunotherapy early before non-restorable cerebellar damage and to maintain adequate intensity and course (16). The similar clinical manifestations with cerebellar ataxia as the main complaint, inflammation of CSF and effectiveness of immunotherapy suggests that anti-EEF1D autoantibodies potentially can be used as biomarker in ACA. The discovery of anti-EEF1D antibodies will contribute to the timely diagnosis of ACA, which hopefully can improve patients’ outcomes.

In summary, we identified an anti-EEF1D antibody as a new autoantibody in two ACA patients. Although the role of this novel antibody in the disease mechanism warrants further study, our findings expand the spectrum of diagnostic anti-neuronal antibodies of ACA.

## 5 Conflict of Interest

The authors declare that the research was conducted in the absence of any commercial or financial relationships that could be construed as a potential conflict of interest.

## 6 Ethics Statement

This study was approved by the Institutional Review Board of Peking Union Medical College Hospital (PUMCH) (JS-891 and JS-2184). Written informed consent was obtained from the patient.

## 7 Author Contributions

HG and JW designed the study and drafted the manuscript. LG, HR, and XC performed the experiments. SF and ML performed clinical examination. All authors have read and approved the final manuscript.

## 8 Funding

This study was funded by National Natural Science Foundation of China (31970648), National High Level Hospital Clinical Research Funding (2022-PUMCH-B-120), Scientific Foundation of Institute of Psychology, Chinese Academy of Sciences (E2CX3815CX). This work was also supported by CAS Key Laboratory of Mental Health, Institute of Psychology, Chinese Academy of Sciences.

## Acknowledgments

We acknowledge funding support from the CAS Key Laboratory of Mental Health, Institute of Psychology, Chinese Academy of Sciences and National Key Research and Development Program of China.

## Notes

### Competing Interest Statement

The authors have declared no competing interest.

